# Respiration competes with theta for modulating parietal cortex neurons

**DOI:** 10.1101/707331

**Authors:** Felix Jung, Yevgenij Yanovsky, Jurij Brankačk, Adriano BL Tort, Andreas Draguhn

**Affiliations:** Institute for Physiology and Pathophysiology, Heidelberg University, 69120 Heidelberg, Germany; Brain Institute, Federal University of Rio Grande do Norte, Natal, RN 59056-450, Brazil

**Author notes:** Equal contribution as senior authors (Adriano BL Tort, Andreas Draguhn). Correspondence (A.B.L. Tort) or (A. Draguhn).

## Abstract

Recent work has shown that nasal respiration entrains local field potential (LFP) and neuronal activity in widespread regions of the brain. This includes non-olfactory regions where respiration-coupled oscillations have been described in different mammals, such as rodents, cats and humans. They may, thus, constitute a global signal aiding interregional communication. Nevertheless, the brain produces other widespread slow rhythms, such as theta oscillations, which also mediate long-range synchronization of neuronal activity. It is completely unknown how these different signals interact to control neuronal network activity. In this work, we characterized respiration- and theta-coupled activity in the posterior parietal cortex of mice. Our results show that respiration-coupled and theta oscillations have different laminar profiles, in which respiration preferentially entrains LFPs and units in more superficial layers, whereas theta modulation does not differ across the parietal cortex. Interestingly, we find that the percentage of theta-modulated units increases in the absence of respiration-coupled oscillations, suggesting that both rhythms compete for modulating parietal cortex neurons. We further show through intracellular recordings that synaptic inhibition is likely to play a role in generating respiration-coupled oscillations at the membrane potential level. Finally, we provide anatomical and electrophysiological evidence of reciprocal monosynaptic connections between the anterior cingulate and posterior parietal cortices, suggesting a possible source of respiration-coupled activity in the parietal cortex.

## Introduction

The brain often produces rhythmic patterns of activity that can be detected at different spatial scales, from intracellular recordings, through local field potential (LFP) and multi-unit activity, up to more macroscopic (EEG, fMRI) measurements [1]. Brain oscillations may thus help to fill the gap between the cellular and network levels and have been suggested to play important roles in brain computations, such as assisting the binding of distributed cell assemblies and the control of information flow [2–5].

In addition to the central nervous system, it is well known that many other physiological systems also exhibit rhythmicity at the ultradian timescale, such as the digestive, cardiovascular and respiratory systems [6–9]. Interestingly, recent studies are starting to unveil relationships between oscillatory activity produced by different organ systems. For instance, Tallon-Baudry and collaborators have shown that ascending signals from the heart and gastrointestinal tract modulate brain dynamics and affect perception, emotion and cognition [10–13]. In particular, the EEG alpha rhythm was shown to be modulated by the phase of the gastric cycle [7]. In a parallel line of work, we and others have recently described that nasal breathing rhythmically modulates neuronal activity in several brain regions [14–26].

Although the first reports of neuronal oscillations coupled to respiration date back to the seminal papers by Adrian [27,28], they were generally believed to occur in olfactory regions [but see 29]. However, a boom of papers published in the last 5 years convincingly demonstrated that rhythmic breathing also entrains non-olfactory networks [e.g., 14–25]. Though respiration-coupled oscillations tend to be most prominent in frontal regions, they can also be detected at smaller magnitudes in more posterior neocortical regions, even in the visual cortex [25]. Such a widespread influence could potentially relate to the effects of respiration over emotional states and cognitive functions [30–34]. Noteworthy, it has been increasingly recognized that nasal respiration modulates performance in cognitive tasks [35–37], including non-olfactory ones [38].

Despite the recent enthusiasm on brain activity coupled to respiration, many questions remain open, constituting an active field of research [26]. To gain further insight into the apparent global influence of nasal breathing, in this work we aimed at studying a distant brain region from the olfactory bulb not primarily related to olfaction. We have thus investigated the posterior parietal cortex, which has been recently shown to exhibit respiration-coupled oscillations at low amplitude [25,39]. In this region prominent theta oscillations can also be detected, thus we paid particular care in differentiating the network effects of theta from respiration. Our results show that both slow rhythms have differential signatures and, interestingly, seem to compete with each other for neuronal entrainment. We also provide evidence that the frontal cortex, which is the neocortical region with largest respiration-coupled oscillations in the rodent [25], could be responsible for mediating respiration-entrained signals into the posterior parietal cortex.

## Results

In order to characterize respiration-coupled brain activity in non-olfactory regions, we chose to investigate the posterior parietal cortex, also known as parietal association cortex (PAC). This region has been shown to exhibit both hippocampal-coherent theta oscillations and local field potential (LFP) rhythms entrained by nasal respiration [25,39], the latter also referred to as the “respiration-rhythm” (RR). The relative magnitude of theta and RR in PAC is highly dynamic and depends on behavior. For instance, RR dominates during immobility while theta oscillations are much more prominent during exploration and REM sleep [25,39]. Here we performed retrograde tracing and electrophysiological recordings in PAC of freely moving and urethane-anesthetized mice. Retrograde tracing aimed at identifying direct afferent projections from the anterior cingulate cortex (ACC), a frontal region that displays prominent RR activity [25]. Field and unit responses to electrical stimulation of the ACC complemented the tracing results. LFPs and neuronal activity were recorded simultaneously with independent measures of respiration (see Materials and Methods).

### Respiration-entrained rhythm in the parietal cortex

LFP recordings confirmed our previous finding that respiration can entrain LFP oscillations in PAC [25,39]. Figure 1A shows example LFPs in the olfactory bulb (OB) and PAC along with the respiration signal (Resp) simultaneously recorded from an awake immobile mouse using whole-body plethysmography. Notice the presence of respiration-coupled LFP oscillations not only in OB, as classically described [27,40,41], but also in PAC (better inferred from the inspection of the filtered traces [light gray lines]). Figure 1B shows power spectra of Resp and LFPs from PAC and OB as well as coherence between PAC and Resp and between PAC and OB. Although RR is much more prominent in the OB (notice the ten-fold difference in power scale between OB and PAC), both the PAC power spectrum and the coherence spectra between PAC and Resp or between PAC and OB exhibit a clear peak at the same frequency as the breathing rate, therefore marking the presence of RR. Of note, RR has been operationally defined as a power peak at the same frequency as and coherent with respiration [26].

**Figure 1.**
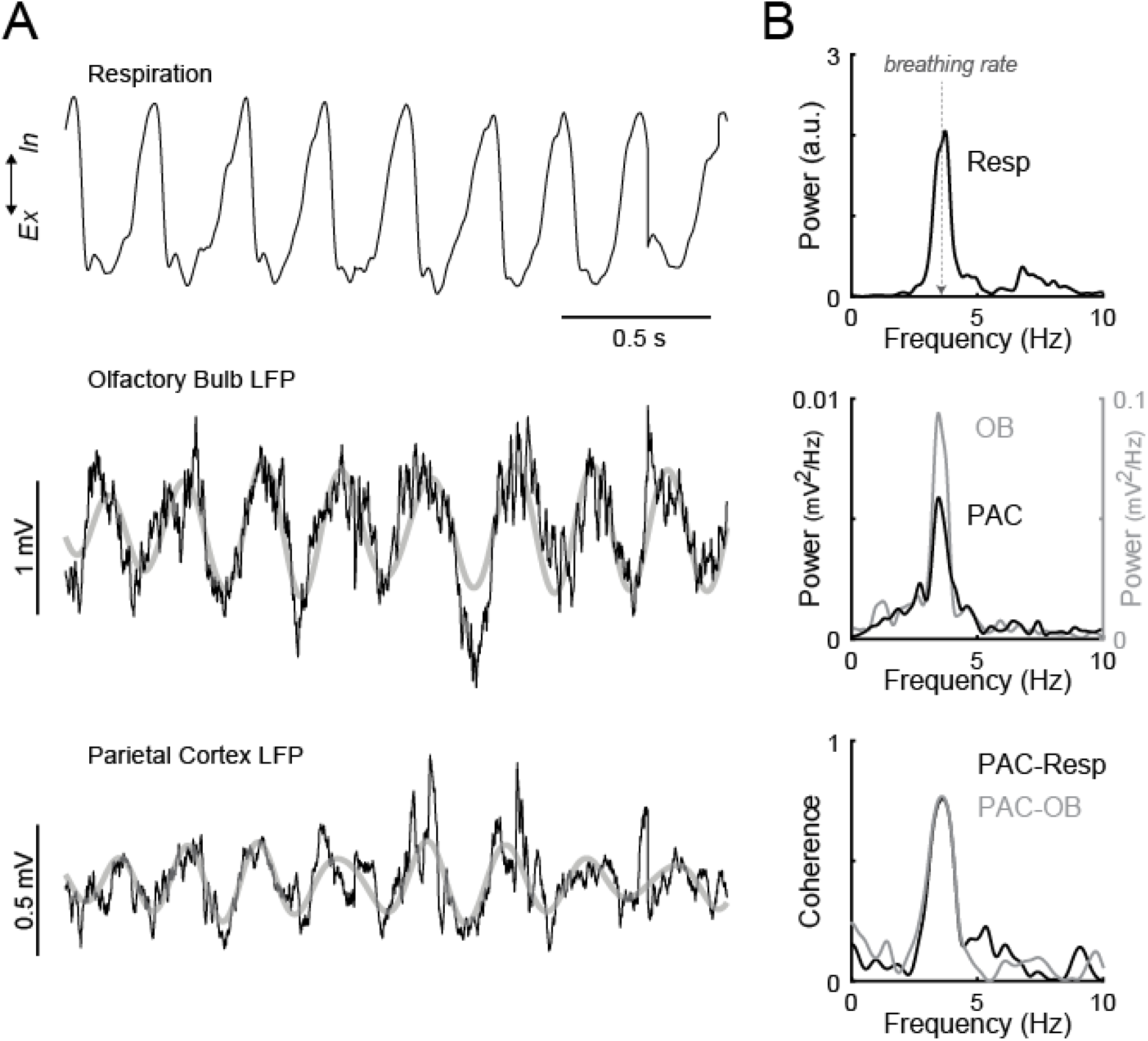
Respiration-coupled LFP oscillations in the posterior parietal cortex (PAC). (A) The top trace shows the respiration signal (Resp) recorded from a mouse through whole-body plethysmography. The bottom traces show OB and PAC LFPs recorded simultaneously (black) along with their band-pass (3-5 Hz) filtered components (gray). (B) Top panels show the power spectra for Resp and LFP signals, as labeled. The bottom panel shows the phase coherence spectrum computed between the PAC and OB LFPs, as well as between PAC LFP and Resp. Notice prominent power and coherence peaks at the same frequency as respiration, which hallmark respiration-coupled oscillations [26]. Data in A-B from the same representative, awake mouse during immobility (epoch length: 20 seconds). Ex: expiration; In: inspiration.

### LFP coherence to respiration decreases with cortical depth

Next, we investigated RR in PAC as a function of recording depth, in the absence of theta oscillations. For this purpose, we implanted 16-channel linear probes with 100 μm electrode distances into the PAC and recorded mice during periods of immobility. The highest channel (channel 1) was placed 100 μm above the cortical surface, and the deepest (channel 16) reached the hippocampal CA1 region below the pyramidal cell layer. Figure 2A shows a typical histological section of an implanted animal. Figure 2B shows an example of simultaneously recorded Resp and LFPs from channel four (200 μm below cortical surface) and channel fourteen (depth 1200 μm, just above hippocampal pyramidal cell layer). The corresponding power and coherence spectra are shown in Figure 2C. Note that the LFP in channel four is much more coherent with respiration than the LFP in channel fourteen. The average heat map of RR activity (Figure 2D; n = 10 mice; two cycles shown) reveals the absence of phase shifts across cortical depths. Next, for each mouse we divided RR amplitude values at each depth by the maximal amplitude across depths (so that the maximal amplitude became 1); Figure 2E depicts the average and individual laminar profiles of normalized RR amplitude. Notice that, on average, RR did not exhibit major amplitude changes across depths. Interestingly, however, we found that the average LFP coherence to respiration decreased with cortical depth and was lowest in CA1 (Figure 2F; n = 10 mice).

**Figure 2.**
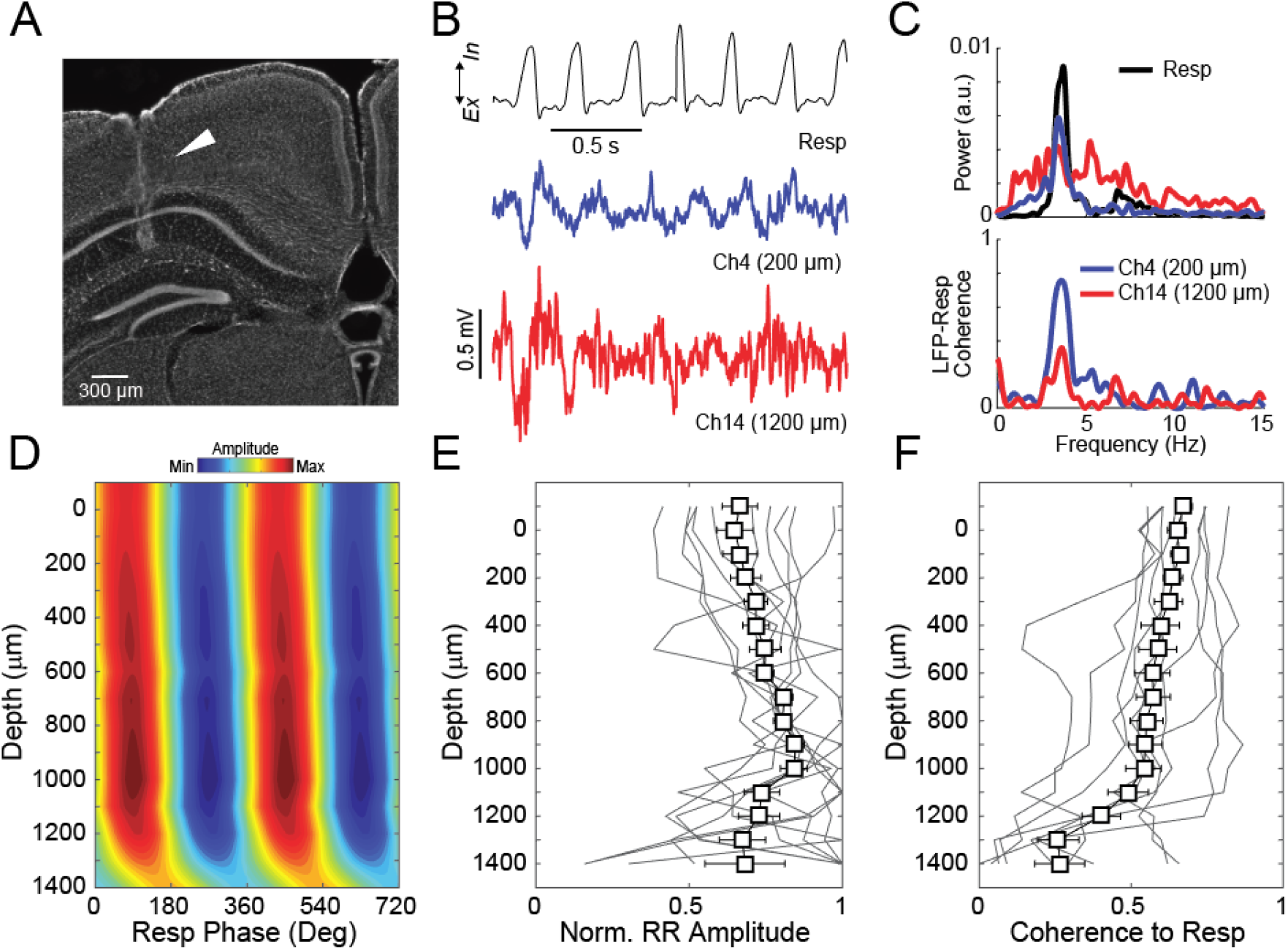
Laminar profile of respiration-coupled LFP oscillations. (A) Histological section showing a linear probe track. (B) Respiration signal (Resp, top) and LFPs (bottom) from two recording locations in an immobile animal. (C) Corresponding power spectra (top) and LFP-Resp coherence (bottom) for each recording location. Notice lower coherence at the deepest location from the brain surface. (D) Amplitude-depth profile of respiration-coupled oscillations (see Methods). (E, F) Normalized RR amplitude (E) and coherence to respiration (F) across recording depths. D-F depict average over 10 mice; errorbars denote SEM; gray lines show data from individual animals. Ex: expiration; In: inspiration; Ch: channel; RR: respiration-entrained LFP rhythm.

### Theta and respiration-coupled oscillations have different laminar profiles

During REM sleep both theta oscillations and RR are simultaneously present in PAC and often separated by frequency (that is, breathing rate is often lower than theta frequency during sleep). Figure 3 compares the laminar profiles of theta and RR during REM sleep in mice implanted with linear multi-contact probes. Figure 3A (upper panel) shows LFPs from channel two (cortical surface) and channel twelve (depth 1000 μm) simultaneously recorded with respiration using whole-body plethysmography. The corresponding power spectra, as well as the coherence spectra between LFPs and Resp, are shown in the lower panel of Figure 3A. Note that theta power (peak between 6 and 10 Hz) is larger at 1000 μm compared to the cortical surface. Notice further that RR power (peak between 3 and 5 Hz, i.e., at the same frequency as the Resp power peak) is much smaller compared to theta power in both locations. Nevertheless, only RR, but not theta, exhibits high coherence to respiration, and moreover, RR-Resp coherence is larger at the surface compared to 1000 μm. Figure 3B depicts the laminar profile of RR and theta, both averaged across ten animals. While the normalized theta amplitude increases with depth, the variations of RR amplitude across depths do not show a clear trend (Figure 3C). Finally, we next calculated the coherence of RR and theta throughout their depth profiles. The first channel in the probe served as reference (note that, unlike in Figure 2F, respiration cannot serve as a reference for this comparison as theta is not coherent to respiration). Interestingly, RR coherence strongly decreases with cortical depth, whereas theta coherence to channel one is stable throughout the cortex (Figure 3D). Thus, we conclude that the laminar profiles of RR and theta activity differ, indicating distinct origins of the two slow rhythms.

**Figure 3.**
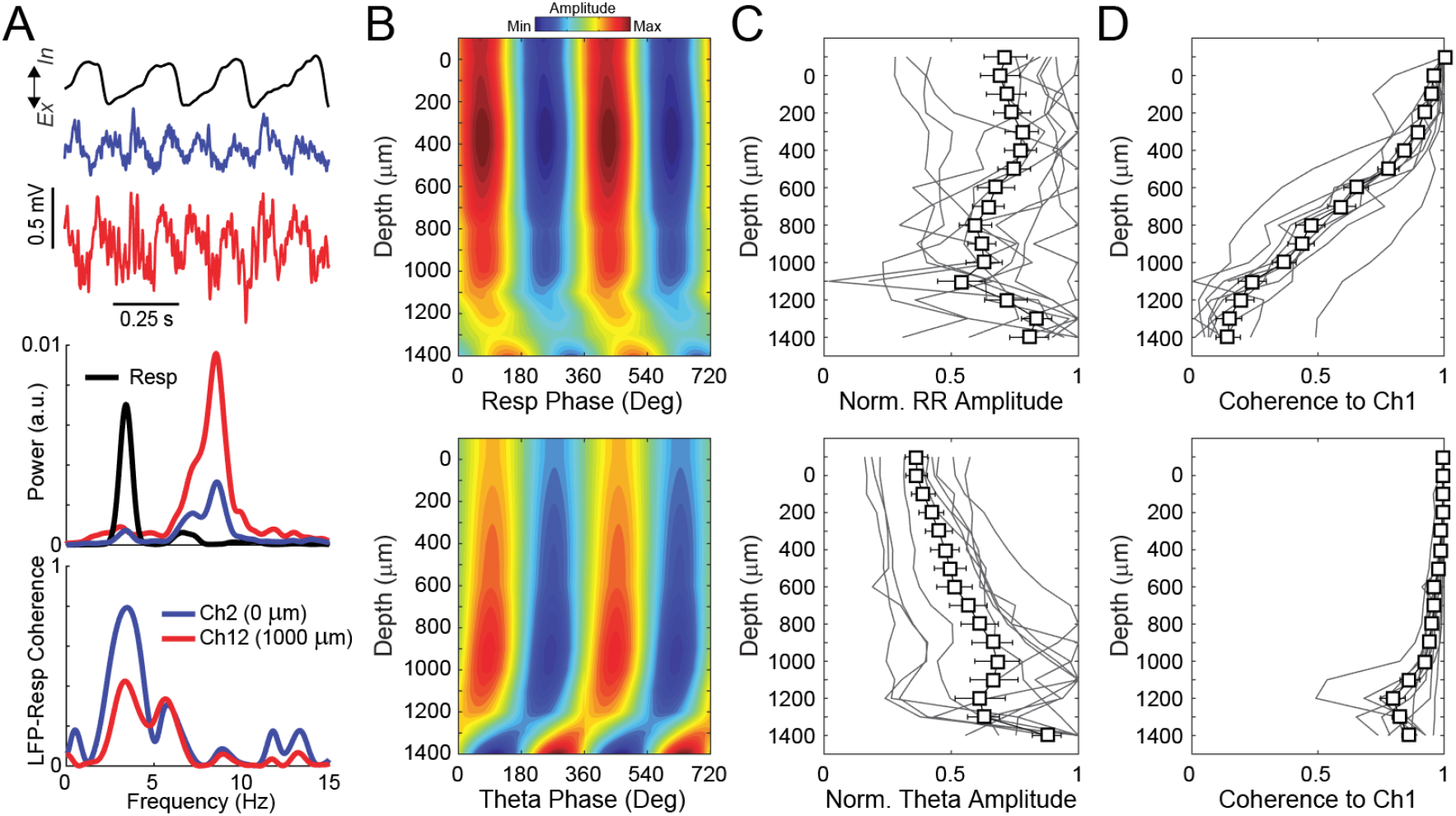
Theta and respiration-coupled oscillations have different laminar profiles. (A) Top traces show respiration and simultaneously recorded LFPs from two cortical depths during REM sleep. The middle panel shows the corresponding power spectra. Notice larger theta power in the deepest channel and similar LFP power levels at the respiration frequency. The bottom panel depicts LFP-Resp coherence for the same channels. Notice that coherence to respiration is lower in the deepest channel. (B) Amplitude-depth profile of RR (top) and theta (bottom). (C) Normalized amplitude of RR (top) and theta (bottom) across recording depths. (D) Coherence to the most superficial channel at respiration (top) and theta (bottom) frequency. B-D depict average over 10 mice; errorbars denote SEM; gray lines show data from individual animals. Ex: expiration; In: inspiration; Ch: channel; RR: respiration-entrained LFP rhythm.

### Respiration competes with theta for modulating parietal cortex neurons

Network oscillations are able, and may actually serve to entrain discharges of individual neurons. We therefore collected juxta- or intracellular recordings of PAC neurons in urethane-anesthetized mice. Typical sleep-like up-and-down activity was suppressed by arousing the animal with light air puffs or tail pinches, evoking simultaneous RR and theta oscillations [15,17]. For the following analysis we only used episodes in which RR and theta could be clearly separated by frequency. Figure 4A shows an example of a PAC neuron recorded juxtacellularly along with Resp and PAC LFP recordings; also shown are the corresponding power spectra and spike-phase distributions. Notice that the LFP power spectrum has a prominent peak between 2 and 4 Hz, matching the Resp power peak and thus corresponding to RR (coherence to Resp is also high [not shown, but see Figure 5B]). The LFP – but not Resp – also exhibits a second, smaller power peak between 4 and 6 Hz, which corresponds to the arousal-induced theta rhythm under urethane anesthesia [15]. Inspection of the spiking probability per respiration phase and per theta phase reveals that this example PAC neuron was modulated by respiration but not theta. Figure 4B shows similar data from a juxtacellularly recorded unit which was modulated by theta but not respiration. A total of 165 PAC units with background LFP activity as defined above were recorded. Sixty-one units were collected in upper layers (above 400 μm) and one hundred and four in lower layers (below 400 μm). Figure 4C (left) shows, separately for the upper and lower layers, the proportion of neurons modulated only by theta, only by respiration, by both rhythms or by neither rhythm. We found that a greater proportion of neurons are modulated by respiration at upper layers compared to lower layers (χ^2^, p<0.05). In contrast, the proportion of theta-modulated units does not differ between the upper and lower layers. Next, we recorded neurons after bypassing nasal respiration through tracheotomy, a procedure previously shown to eliminate RR while maintaining theta activity [15,17]. A total of sixty-six neurons were recorded in urethane-anesthetized animals after tracheotomy, twenty-five above 400 μm and forty-one below 400 μm. Under this protocol, the proportion of neurons modulated by theta significantly increased in relation to nasal breathing at both recording locations (Figure 4C right). Finally, Figure 4D shows the coupling strength of spikes to the phase of either Resp or theta (before and after tracheotomy), plotted as a function of the mean spiking phase. Interestingly, the distribution of the mean spiking phase within theta cycles differs between nasal and tracheal breathing (Kuiper test, p<0.05). Together, these results show that respiration differentially modulates units at upper and lower cortical layers, and that, in the absence of RR (induced by tracheotomy), theta modulates more units than during nasal breathing. The latter finding indicates that both slow rhythms compete for controlling neuronal activity.

**Figure 4.**
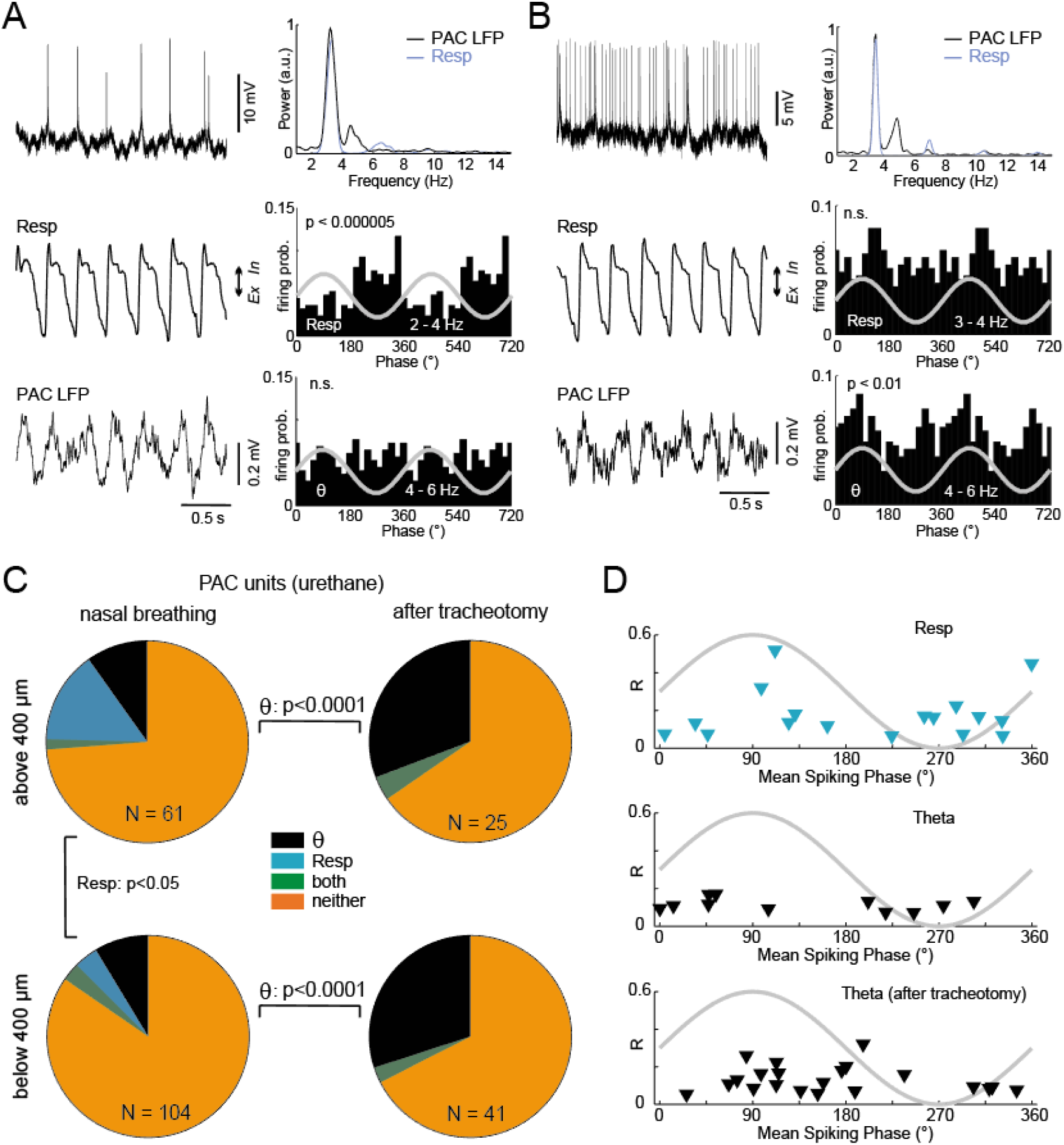
Respiration competes with theta for modulating parietal cortex neurons. (A) Left: traces show a juxtacellular recording of a PAC neuron (top), along with the respiration (Resp, middle) and LFP (bottom) signals recorded simultaneously in a urethane-anesthetized mouse. Right: the top panel shows power spectra for Resp and PAC LFP. Notice in the LFP spectrum a power peak at the same frequency as respiration and a second power peak corresponding to urethane-induced theta (θ) activity. The middle panel shows spiking probability per respiration phase and the bottom panel per theta phase. Notice that this example neuron was modulated by respiration but not theta. (B) Same as in A, but for a juxtacellularly recorded unit modulated by theta only. (C) Left: proportion of neurons modulated only by theta, only by respiration, by both rhythms or by neither rhythm, shown separately for recording locations above and below 400 μm from cortical surface. Respiration modulates a greater proportion of neurons at more superficial recording sites (χ^2^, p<0.05), while the proportion of units modulated by theta is not significantly different. Right: same as before, but for neurons recorded after tracheotomy. The proportion of units modulated by theta significantly increases at both superficial and deep locations. (D) Spike-phase coupling strength (R) vs the mean spiking phase for Resp (top), and theta before (middle) and after (bottom) tracheotomy. Ex: expiration; In: inspiration.

**Figure 5.**
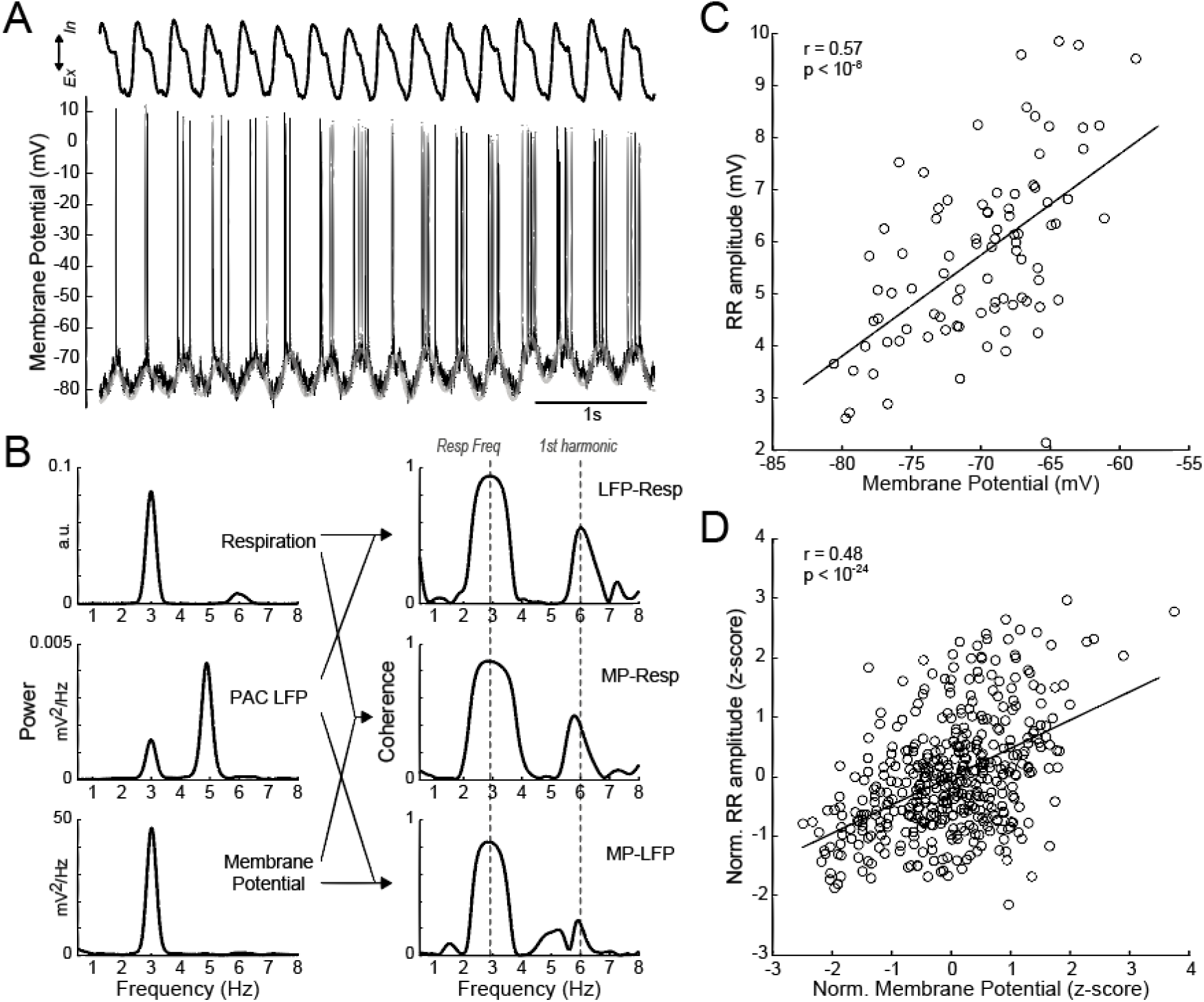
Respiration-coupled oscillations can be detected intracellularly and are likely to depend on GABAergic synapses. (A) Membrane potential (MP) of a PAC neuron simultaneously recorded with respiration in a urethane-anesthetized mouse (recording depth: 269 μm). Superimposed gray trace shows the low-pass (<10 Hz) filtered MP. Notice MP oscillations coupled to breathing cycles. (B) Left panels show power spectra for MP, PAC LFP and Resp. Notice that the MP exhibits only one power peak, which corresponds to RR, while the LFP also exhibits a second power peak which corresponds to theta. The right panels show phase coherence spectra for all pairwise combinations. Notice in all cases power and coherence peaks at the same frequency as respiration. (C) Scatter plot of intracellular RR amplitude vs MP for the same example cell. Notice reduction of RR amplitude at more hyperpolarized potentials. (D) Pooled scatter plot of RR amplitude vs MP, expressed as z-score units (data from 6 PAC neurons exhibiting intracellular RR). Ex: expiration; In: inspiration; RR: respiration-entrained MP rhythm.

### Respiration-coupled oscillations can be detected intracellularly and likely depend on GABAergic synapses

In order to gain insight into the subthreshold signals mediating entrainment by RR or theta, we next performed intracellular recordings from PAC neurons in urethane-anesthetized mice. We subjected these animals to arousal stimuli (tail pinch or air puffs), so as to elicit theta oscillations [15]. Noteworthy, from a total of twenty-nine cells that could be successfully recorded, six PAC neurons exhibited clear membrane potential (MP) oscillations coupled to respiration (4 cells above 400 μm, 2 cells below). Figure 5A shows the MP of one such PAC neuron along with the simultaneously recorded respiration signal. The low-pass filtered MP, which exhibits no action potentials, is shown superimposed. Notice prominent MP oscillations phase-locked to breathing cycles. The power spectra of MP and Resp, as well as the power spectrum of the PAC LFP and the coherence spectra between all pairwise combinations of these signals, are shown in Figure 5B. Interestingly, the MP exhibits only one power peak, which corresponds to RR, whereas the LFP also exhibits a second peak corresponding to theta (notice further that theta has larger power than RR in the LFP). Both the MP and the PAC LFP exhibited coherence peaks with Resp at the same frequency as breathing and at its harmonics (as in non-anesthetized mice [Figure 3], the LFP was not coherent with Resp at theta frequency). Next, we investigated whether the amplitude of respiration-coupled oscillations recorded intracellularly depended on the level of subthreshold depolarization/hyperpolarization of the MP (see Material and Methods). For this same example cell, we found a significant positive linear correlation (r = 0.57, p < 10^−8^) between the amplitude of RR recorded intracellularly and the MP level (Figure 5C). Indeed, for each of the four cells with clear subthreshold RR activity, we found that the amplitude of respiration-entrained MP oscillations increased upon depolarization. This linear correlation could be observed for the pooled scatter plot of RR amplitudes versus MP using data from the four neurons when the variables are expressed as z-score units (r = 0.48, p < 10^−24^; Figure 5D; the z-score normalization was necessary because of differences in MP levels across neurons). The increase of RR amplitude with depolarizing MP above −80 mV suggests involvement of chloride channels; therefore, the respiratory signal is likely to modulate the PAC neurons via GABA-A receptor-mediated inhibition. Interestingly, from 22 cells that were intracellularly recorded while the PAC LFP exhibited theta oscillations during normal breathing, only one cell had theta at the MP level (this cell also had intracellular RR), while none of 7 cells recorded during tracheal breathing in the presence of LFP theta exhibited theta intracellularly.

### Anterior cingulate cortex may be a source of respiration-coupled activity in the parietal cortex

Next, we wondered what could be the upstream source for the respiration-coupled activity in PAC. A possible candidate region is the anterior cingulate cortex (ACC), a frontal area in which we recently found prominent RR [25] and, moreover, which has been previously reported to connect with the posterior parietal cortex [42–45]. We first sought to corroborate the existence of direct projections from ACC to PAC in mice. To that end, we injected rabies virus or retrobeads into the PAC of six mice to identify monosynaptic afferents by retrograde tracing. Figure 6A depicts the injection site in PAC (left panel) along with neurons in the frontal cortex (AP +0.14 mm) retrogradely labeled by rabies virus (red dots in the middle panel) and by retrobeads (green dots in the right panel). Both labeling methods demonstrate monosynaptic afferents from frontal regions including the ACC (labeled “Cg1” in the middle and right panels). In a second approach, we used multi-contact linear probes to record laminar profiles of responses evoked by electrical stimulation of ACC. Figure 6B shows averaged evoked responses across PAC following electrical stimulation of the ACC in awake immobile animals (n = 6 mice). ACC stimulation led to an earlier transient response at more superficial recording sites in PAC followed by a slower component in deeper sites.

**Figure 6.**
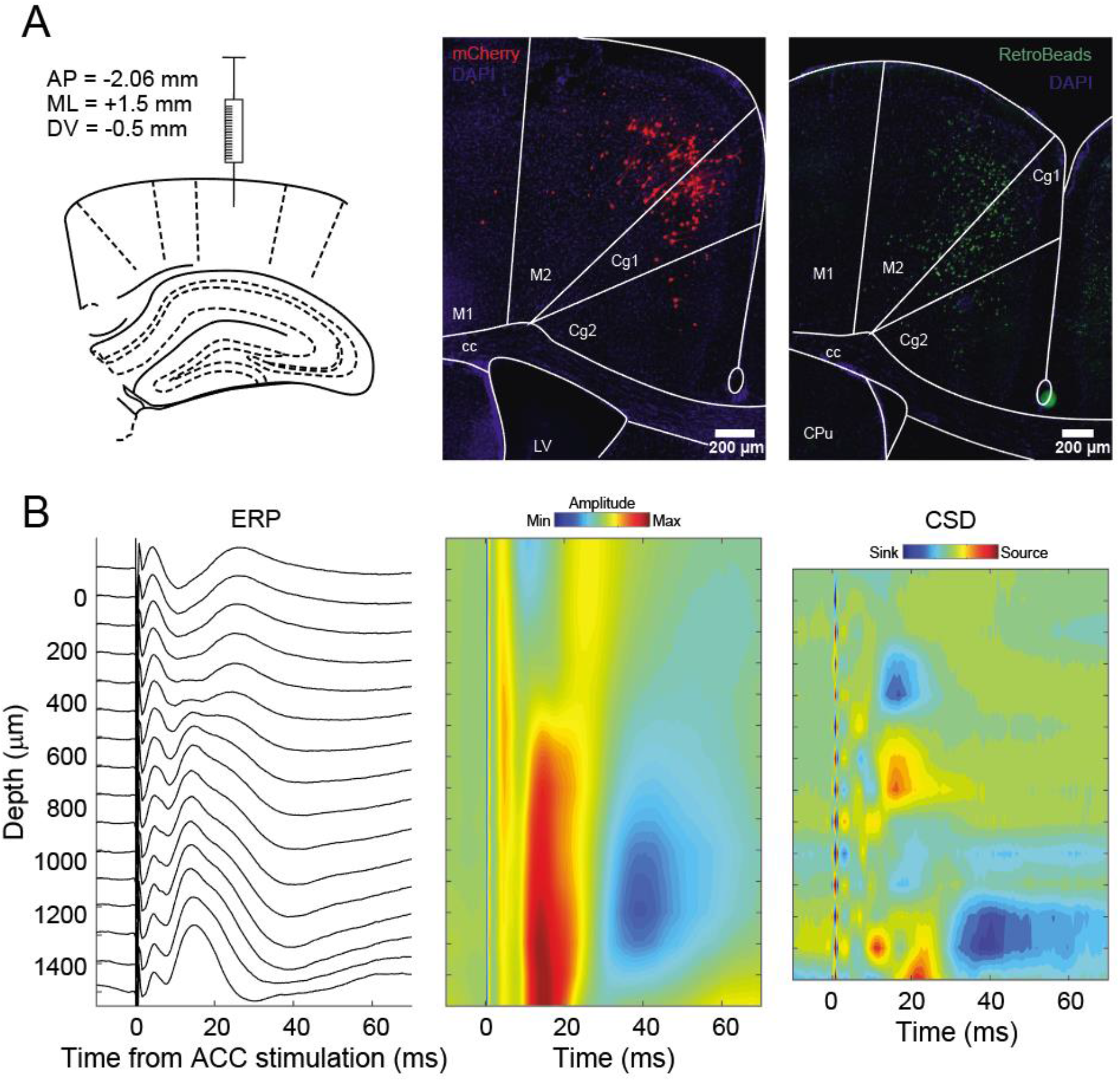
Anterior cingulate cortex may be a source of respiration-coupled activity in the parietal cortex. (A) Left: schematic depicting the injection of rabies virus or retrobeads into the posterior parietal cortex. The middle and right panels show retrogradely labeled neurons in frontal cortex (AP: +0.14 mm) through either method (middle: rabies virus; right: retrobeads). Notice labeled neurons in the cingulate cortex (Cg1), which has been previously shown to exhibit prominent respiration-coupled oscillations [25]. (B) Left traces show average evoked response potentials (ERP) across the parietal cortex following electrical stimulation of the anterior cingulate cortex (ACC) in awake immobile mice. The middle and right panels show the average ERP and associated average current source density (CSD) in pseudocolor scale (n=6 mice). Notice an earlier fast component at more superficial sites followed by a slower response in deeper sites. AP: anteroposterior; ML: mediolateral; DV: dorsoventral.

### The anterior cingulate cortex modulates spiking activity in the parietal cortex

In addition to the laminar profiles of awake mice (Figure 6), we also recorded spiking activity in response to electrical stimulation of the ACC in urethane-anesthetized mice. Figure 7A shows a typical peristimulus time histogram from a single PAC neuron to 27 ACC stimulations as well as the MP response of this cell to a single stimulation. Both reveal a short-latency excitation with the emission of single spikes followed by a ~200 ms inhibition of spiking activity. Interestingly, in addition to orthodromic responses, we also found antidromic spikes followed by EPSPs and IPSPs (Figure 7B) exclusively in deep cortical laminae (red dots in Figure 7B, right panel). This observation points to a reciprocal monosynaptic connection between PAC and ACC. In contrast to the predominantly deep location of neurons with antidromic responses, orthodromic spike latencies to ACC stimulation were found more evenly distributed in PAC (black dots in Figure 7B, right panel). In addition to orthodromic or antidromic single spikes followed by inhibition, we also found cases of bursts of spikes without inhibition and of IPSPs without previous spikes (Figure 7C). Notice that the latter could be a possible source of intracellular RR (Figure 5).

**Figure 7.**
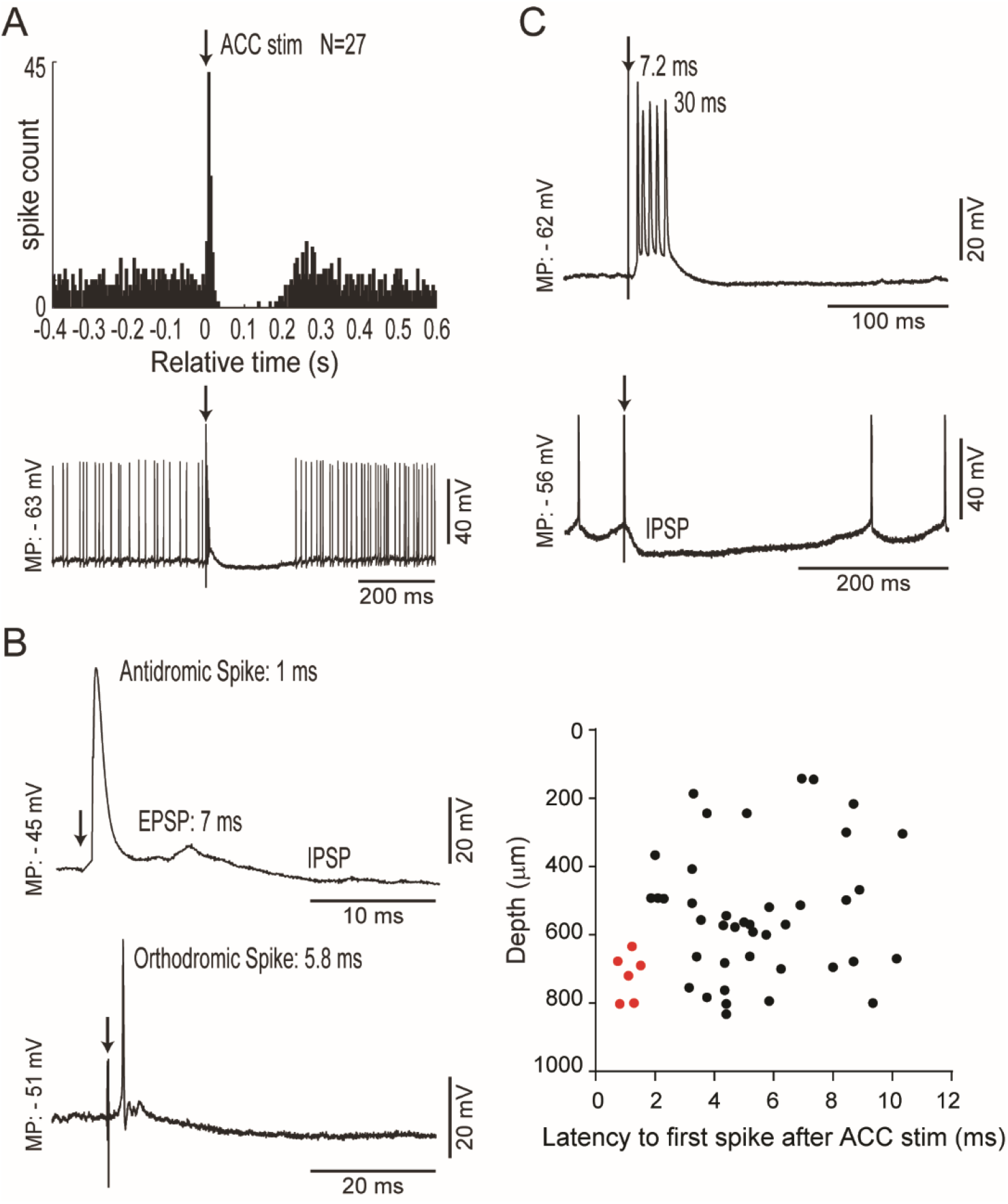
The anterior cingulate cortex modulates spiking activity in the parietal cortex. (A) Top: Peristimulus time histogram for a single neuron in the parietal cortex (PAC) relative to electrical stimulation (down arrow) of the anterior cingulate cortex (ACC). Notice that ACC stimulation leads to a sharp increase in neuronal activity followed by suppression. The bottom trace shows the intracellular recording of this example PAC neuron during one stimulation trial. (B) The left traces show examples of an antidromic (top) and an orthodromic (bottom) spike in PAC following ACC stimulation (down arrow). The right panel depicts the latency to the first spike following ACC stimulation according to the recording depth. Putative antidromic spikes (latency < 2 ms, shown in red) were only found in deep PAC layers. (C) Example of a PAC neuron highly excited (top) and a PAC neuron highly inhibited (bottom) following ACC stimulation (down arrow).

## Discussion

Brain oscillations are far from being the only temporal rhythmic patterns produced by the body; in fact, many physiological systems work rhythmically. Among them, the digestive, cardiovascular and respiratory systems are well known to exhibit rhythmicity at the ultradian timescale [6–9]. Recent research has started to unveil links between the brain and other body rhythms [12,26]. In particular, the coupling between neuronal activity and the cycles of nasal breathing has been increasingly recognized [14,15,19]. Although such effect has been long described for olfactory regions [27,28], the novelty resides in the discovery that respiration also modulates many other brain areas which are typically not associated with odor processing [16,21,24,25].

In this work, we confirm that rhythmic breathing modulates the activity of an associative cortical area not primarily involved with olfaction. Noteworthy, we found respiration-coupled activity at multiple spatial scales in the posterior parietal cortex: from the mesoscopic, network level (as inferred by LFP analysis), passing through the level of action potentials (as inferred by spike-phase coupling), and down to the subthreshold membrane potential level (as inferred by intracellular recordings). It should be noted, however, that even though our results show that respiration does impact parietal cortex activity, the amplitude of RR is rather low in this region. Indeed, we have recently shown that respiration-coupled activity has an anteroposterior gradient in mice, in which it tends to be larger in the frontal regions [25]. Nevertheless, by concomitantly recording respiration and employing careful signal analysis, RR can be clearly detected in more posterior regions [25,39]. Due to its small amplitude, however, RR is easily missed when it occurs along with the dominant theta rhythm in awake animals (e.g., Figure 2). Therefore, in the absence of concurrent tracking of respiration, RR is likely to remain unnoticed in this region. However, the Resp-LFP phase coherence spectrum, which can only be performed when tracking respiration, clearly reveals that a component of the LFP spectrum is phase-locked to breathing cycles and hence RR entrains network activity in the parietal cortex. Moreover, the tracking of respiration further allows inferring the existence of PAC neurons modulated by breathing cycles, despite the low amplitude of RR in PAC.

A common issue in LFP analysis is to infer whether the detected oscillations represent local activity (presumably synaptic) or else passive/return currents due to volume conduction from current dipoles located elsewhere [1,46,47]. Although RR has low amplitude in the parietal cortex, our results support the notion that it does directly and specifically impact the local network, which in turn is likely to contribute to RR appearance at the LFP level. In particular, we found that RR modulates spike timing, which is considered an unlikely effect of volume conduction [but see 48,49]. Moreover, we found that RR modulation of neurons depends on recording depth: more units couple to respiration in the superficial than deep layers, an effect that we again deem unlikely to be due to volume conduction but instead point towards specific connectivity patterns. Most convincing evidence comes from intracellular recordings which revealed clear cases of subthreshold membrane potential oscillations phase-locked to breathing cycles (Figure 5). Interestingly, in all these cases the amplitude of intracellular RR decreased with more hyperpolarized membrane potentials, indicating involvement of GABAergic transmission. This fits well with our results from responses to stimulation of the anterior cingulate cortex which induced inhibitory periods of about 200 ms duration (Figure 7A). This time window corresponds to a frequency of 5 Hz, i.e. close to the frequency defined by respiration.

Respiration and theta frequency may often overlap, making it challenging to disentangle both rhythms through standard signal analysis techniques [25,26]. Therefore, we have here focused on periods in which breathing rate was slower than theta frequency, so that, by band-pass filtering, we could separately investigate these rhythms. Under this approach, we found that respiration entrainment and theta have different laminar profiles across the parietal cortex-hippocampus axis. Namely, at theta frequency, LFP coherence to the most superficial electrode was stable across recording depths (Figure 3), whereas, at the breathing frequency, LFP coherence to the first electrode in the probe, as well as to the respiration signal itself, decreased for deeper locations (Figures 2 and 3). Regarding amplitude, theta oscillations became larger with depth, especially closer to the hippocampus, while RR amplitude did not exhibit a consistent trend across animals. Most strikingly, in addition to the LFPs, we found that respiration preferentially entrained units in more superficial layers (Figure 4). In fact, respiration modulated a greater percentage of units (16.4%) than theta (11.5%) in the superficial layers, even though RR has lower amplitude than theta at the LFP level. Interestingly, we further note that the percentage of neurons modulated by respiration in deep layers (6.7%) was not much greater than the false positive rate determined by the significance threshold (i.e., α = 5%). In contrast to respiration, theta modulation of neurons did not differ across the parietal cortex (11.5% of neurons in both superficial and deep layers). In all, these observations corroborate that theta and RR have independent mechanisms of generation.

It has been previously shown that RR depends on nasal airflow, and, as such, it is abolished when animals breath straight through the trachea [15,17]. Interestingly, in addition to RR absence, the proportion of parietal cortex neurons modulated by theta roughly tripled when animals breathed through a tracheostomy (11.5% vs 33.3%, p<0.05, χ2). This suggests that theta and respiration compete with one another for controlling neuronal activity. The lower proportion of neurons modulated by theta in the presence of respiration-coupled oscillations could be due to some neurons having weaker inputs associated with theta than respiration. In this scenario, respiration-coupled postsynaptic potentials would have a larger amplitude than theta-paced inputs, and thus theta modulation would only become noticeable in the absence of respiration-coupled synaptic inputs. Another possibility is a competitive effect at the network level, in which the absence of RR would lead to disinhibition of neurons mediating theta effects.

We note, however, that, except for one cell, we did not detect MP theta activity at the subthreshold level in all other posterior parietal cortex neurons that were intracellularly recorded in the presence of LFP theta oscillations (n = 21 cells), even during tracheal breathing (additional 7 cells) when RR was not present. In particular, notice in Figure 5 that despite the LFP exhibiting prominent theta oscillations at a larger amplitude than RR, there is no theta activity inside the neuron. The posterior parietal cortex does not receive direct connections from the hippocampus [45]. Therefore, if this region receives theta-paced inputs, they are likely to be transmitted through the retrosplenial cortex [see 50], which heavily connects to it [45,51]. Whether theta exerts any far field effect in this region is yet to be determined. In any case, it remains puzzling that we could observe extracellularly recorded parietal cortex neurons modulated by the LFP theta but not much evidence of theta activity at the intracellular level. Consistent with this, no intracellularly recorded neuron exhibited spikes coupled to theta. In contrast, spikes emitted by neurons exhibiting intracellular RR were modulated by respiration (see Figure 5A for an example). One possible, technical explanation is that intracellular recordings are mainly successful in a subpopulation of larger cells that would differ from the cell types modulated by theta. Noteworthy, our findings showing that theta modulates extracellularly recorded posterior parietal cortex neurons match those of Sirota et al. [52]. These authors have also reported intracellular oscillations in S1 neurons during urethane anesthesia (they did not record posterior parietal neurons intracellularly), which they ascribed to theta oscillations [52]. Based on our observations, however, it is feasible that the reported intracellular oscillations in S1 could potentially correspond to RR, but this cannot be inferred without concomitant recording of respiration.

Finally, we set out to investigate the anterior cingulate cortex as a possible contributor to RR activity in the posterior parietal cortex. We studied this region because it displays prominent RR [25] and sends projections to a myriad of cortical and subcortical structures [44,53]. Here, we could corroborate that the ACC directly projects to PAC [42–45,51]. Moreover, electrophysiological data showed antidromic spikes in PAC after ACC stimulation (Figure 7), indicating that ACC receives monosynaptic connections from PAC. The latter result is consistent with recent interconnectivity studies in mice showing that ACC and PAC mutually connect [44,51,53]. Of note, one such connectome study proposed that ACC and PAC form part of a same medial subnetwork, which would be responsible for mediating information from sensory and higher-order areas [44]. Accordingly, the posterior parietal cortex is considered an association region since it receives inputs from visual, somatosensory and auditory areas [43,45]. Our results suggest that the integration of the multimodal information arriving in this region may be influenced by respiration-coupled inputs mediated by the ACC. In turn, respiration inputs are likely to arrive in frontal cortical regions via connections from the piriform and insular cortices [44,54].

In summary, we report that LFPs and single neurons in the PAC are modulated by respiration, that theta and RR are generated by different mechanisms, including a role for GABAergic inhibition in RR, and that both rhythms compete for modulating PAC. We also demonstrate that reciprocal monosynaptic connections between ACC and PAC are a possible source of respiratory influence over PAC. Our findings indicate a strong influence of oscillating body signals on brain function, far beyond specific sensory or motor activity. Specifically, they show that neuronal activity in a cortical association area is modulated by both, respiration as well as theta, which entrain network function in a differential and competitive way.

## Material and Methods

Electrophysiological recordings were performed in 46 adult mice of either sex. In ten freely moving mice, we recorded local field potentials (LFPs) in the posterior parietal cortex (PAC) using chronically implanted NeuroNexus silicon probes (16 contacts separated by 100 μm) during two spontaneous network oscillations: theta (θ) and the respiration-entrained rhythm [RR; 15,18]. In six of these animals, we also recorded PAC LFPs during electrical stimulations of the anterior cingulate cortex (ACC). In a further set of experiments, we recorded from PAC neurons in 36 urethane-anesthetized mice to investigate unit modulation by theta and RR. Six of the urethane-anesthetized mice were tracheotomized before the recording session to abolish RR [15,17]. In eleven of the anesthetized mice, we recorded responses of PAC neurons to electrical stimulation of the ACC. In addition, six mice were injected with rabies virus or retrobeads into the PAC to identify monosynaptic afferents from the ACC by retrograde tracing. Below we describe the experiments in detail.

### Ethics statement

Experiments in this study were performed in accordance with guidelines of the European Science Foundation (2001) and the U.S. National Institutes of Health Guide for the Care and Use of Laboratory Animals and have been approved by the Governmental Supervisory Panel on Animal Experiments of Baden Württemberg at Karlsruhe (35–9185.81/G-84/13 for electrophysiology and 35–9185.81/G-194/17 for tracing). All efforts were made to minimize animal suffering and to reduce the number of animals used. Because of the system level approach of our study, alternatives to *in vivo* techniques were not available.

### Animal care and housing conditions

Mice (C57BL/6N) were purchased at 42 or 84 days of age from Charles River. Animals were housed in groups of four inside a ventilated Scantainer (Scanbur BK, Denmark) on an inverted 12/12 h light/dark cycle (light on 8:00 P.M.) for a minimum of 2 weeks (except for the tracing experiments). Animals had free access to water and food. After chronic electrode implantation or tracer injection, mice were housed individually throughout the experiment. Chronic animals were killed with an overdose of ketamine-xylazine during brain perfusion and animals used in acute experiments with an overdose of ketamine-xylazine after the recording.

### Animal preparation and experimental procedures

#### Retrograde tracing

Four male and two female mice (42-70 days old, weight: 20-22 g) were used for retrograde tracing of monosynaptic afferents to PAC. The animals were anesthetized (see below for surgery details), and two different retrograde tracers were used in two different groups of animals. In both groups, tracers were injected into the right PAC (2.06 mm posterior to Bregma, 1.5 mm lateral to the midline, 0.5 mm ventral from the brain surface) with an estimated rate of 50 nl/min using glass micropipettes (Blaubrand intra MARK, Brand GmbH, Germany). The capillary was held in place for 5 min prior and 10 min after the injection before being slowly retracted from the brain. In the first group of three male mice, we injected 300 nl of green fluorophore-coated latex microspheres (RetroBeads^TM^; Lumafluor Inc., Durham, NC, USA) diluted 1:1 in phosphate buffered saline (Katz and Iarovici, 1990). These animals were killed 10-14 days after the injection for further histological processing (see below for details). In a second group of one male and two female animals, we used a rabies virus-based approach of retrograde tracing consisting of two injections separated by 3 weeks [55]. The first injection (300 nl) was composed of AAV1/2-Syn-iCre (kindly provided by Dr. Thomas Kuner, Heidelberg, Germany) and (1:1) AAV1-EF1α-FLEX-GTB (1.08*10^11^ GC/ml; Salk Institute, La Jolla, CA, USA). The second injection contained 800 nl of the rabies virus Rb-EnvA-ΔG-mCherry (3*10^8^ IFU/ml; kindly provided by Dr. Karl-Klaus Conzelmann, Munich, Germany). These animals were killed 7 days after the rabies virus injection. After removing the brain (see below for details) coronal sections of 50 μm were cut, counterstained with 4,6-Diamidin-2-phenylindol (DAPI; Carl-Roth GmbH and Co. KG, Karlsruhe, Germany) and imaged with a fluorescent microscope and an F-View II camera (Olympus K.K., Tokyo, Japan).

#### Chronic electrode implantation

A total of ten male C57BL/6N mice weighing 24–30 g (84–126 days old) were anesthetized with isoflurane in medical oxygen (4% isoflurane for induction, 1.5%-2.5% for maintenance, flow rate: 1 l per minute). For analgesia, 0.1 mg/kg of buprenorphine was injected subcutaneously before and four and eight hours after surgery. Anesthetized animals were mounted on a stereotaxic apparatus (Kopf Instruments) with a custom-made inhalation tube. Body temperature was maintained at 38°C by a heating pad (ATC-2000, World Precision Instruments). After exposure of the skull, holes of 0.5–1.0 mm in diameter were drilled above the right and left PAC, right and left OB and the cerebellum according to stereotaxic coordinates [based on 56]. Two stainless steel watch screws (1 mm diameter, 3 mm length) over the cerebellum served as ground and reference electrode. A third screw was placed at the surface of left PAC (2.06 posterior to bregma, 1.5 mm lateral). Pairs of varnish-insulated tungsten wires (50 μm, glued together) were implanted into the granular layer of left and right olfactory bulb (OB, 4.55 anterior to bregma, 0.8 mm lateral, 1.45 mm ventral). In six animals an additional pair of varnish-insulated nickel chrome wires cut at an angle of 45° was implanted into the right ACC (0.14 mm anterior to bregma, 0.35 mm lateral, 1.25 mm ventral) for electrical stimulation (0.1 ms duration, 150 to 225 μA intensity). Silicon probes (A1×16–3 mm-100–703-CM16LP, NeuroNexus Technologies) were chronically implanted into the right PAC (2.06 posterior to bregma, 1.5 mm lateral, the uppermost electrode was located 100 μm above the cortical surface and the deepest below the CA1 pyramidal cell layer).

#### Electrophysiology in freely moving mice

One week after surgery, recordings began with a 1 h session in the animal’s home cage. For reliable recording of respiration we used whole-body plethysmography [EMKA Technologies, S.A.S., France, for details see 57]. In both the home cage and plethysmograph, movements were detected with 3-D accelerometry. We aimed at collecting periods with RR in the absence of theta, as present during immobility, and with the simultaneous presence of RR and theta, as present during REM sleep [25,39]. For the latter, we looked for periods with non-overlapping frequencies. Therefore, several recording sessions of up to 4 h were performed in the whole-body plethysmograph on consecutive days. Extracellular signals were filtered (1–500 Hz), amplified (RHD2132, Intan Technologies), digitized (2.5 kHz), and stored for offline analyses with custom-written MATLAB routines (The Mathworks, Inc.).

#### Behavioral staging

After visual inspection, artifact-free periods were behaviorally staged. Classification of vigilance states was based on (i) the level of accelerometer activity (active waking > awake immobility, NREM, REM), (ii) the amount of high-amplitude slow-wave activity in the neocortex (NREM > waking, REM), and (iii) the amount of regular theta oscillations in PAC overlaying the dorsal hippocampus (active waking, REM > awake immobility, none in NREM). Awake immobility was defined as the absence of any prominent accelerometer signal and any slow-wave (delta) activity indicating sleep. A detailed description of behavioral staging is provided elsewhere [39,57,58].

#### Urethane experiments

Twenty-five male and eleven female C57BL/6N mice weighing 19–33 g (77–180 days old) were anesthetized with a mixture of urethane (1.2 g/kg) and ketamine-xylazine (10 mg/kg, 1 mg/kg, i.p.). Urethane and ketamine-xylazine were freshly dissolved in isotonic (0.9%) NaCl solution. All solutions were heated to 38°C before application. The level of anesthesia was maintained so that hind limb pinching produced no reflex movement. Supplemental doses of urethane (0.2 g/kg) were delivered as needed (approximately every 1.5 h). The animals were mounted on a stereotaxic frame (Kopf Instruments) and body temperature was maintained at 38°C. After exposure of the skull, holes of 0.5–1.0 mm in diameter were drilled above the right and left PAC according to stereotaxic coordinates [56]. The dura mater was carefully removed and a 125 μm tungsten electrode (MicroProbes) was implanted into the left PAC (2.0 mm posterior to bregma, 1.5 mm lateral, 0.7 mm ventral) to record LFPs. In some animals, PAC LFPs were recorded ipsilaterally at the right PAC (1.96 mm posterior to bregma, 0.5 mm lateral, 0.75 mm ventral) under an angle of 30° to prevent interference with microelectrodes (see below). A pair of 125 μm tungsten electrodes (MicroProbes) glued together cut at an angle of 45° was positioned into the right ACC (0.14 anterior to bregma, 0.35 lateral, 1.25 mm ventral for the deepest electrode) to stimulate ipsilateral afferents to PAC. Respiration was monitored by recording the chest wall movements using a piezoelectric device (EPZ-20MS64, EKULIT) located beneath the animal’s body [15]. Tracheotomy was performed under additional local lidocaine anesthesia previous to recording in six male mice. Extracellular signals from left PAC were filtered (1–500 Hz), amplified (EXT-16DX or EXT 10-2F, npi, Tamm), digitized (20 kHz) using the CED 1401 board, and stored for offline analyses. Intracellular recordings of right PAC neurons were obtained with high-impedance quartz-glass micropipettes (o.d.1.0/i.d.0.5 mm; impedance: 60–120 MΩ) pulled with the P-2000 puller (Sutter Instruments) penetrating the brain under an angle of 15° (2.0 posterior to bregma, 1.5 mm lateral). Recording electrodes were filled with 1 M potassium acetate and slowly lowered in 5 μm steps using the Micropositioner 2660 (Kopf Instruments). Axoclamp 900A (Molecular Devices) in current-clamp mode with a bridge circuit was used to amplify intracellular signals. For unit recordings, signals were filtered (0–10 kHz) and digitized at 20 kHz. Juxta-cellular recordings were performed by using quartz-glass micropipettes with impedance lowered to 10–20 MΩ. Extracellularly recorded multiunit activity (MUA) from low impedance pipettes was obtained by high-pass filtering (>500 Hz). Due to the typically prevailing slow wave-like sleep activity during urethane anesthesia, tail pinches or air puffs were used to suppress electrophysiological sleep activity and induce theta rhythm and RR. This effect had to be carefully controlled since too strong arousal increased the respiration rate and caused a frequency overlap of RR and theta, whereas weak arousal would induce only RR but not theta. Intra- and juxtacellular responses to ACC stimulation (0.1 ms duration, 250 to 400 μA intensity) were recorded during both sleep-like activity and in presence of theta and RR after arousal.

### Data analysis

Data were analyzed in MATLAB (The MathWorks) using built-in and custom-written routines.

#### Spectral analysis and coherence analysis

Power spectral density was calculated by means of the Welch periodogram method using the *pwelch.m* function from the Signal Processing Toolbox (50% overlapping, 4 s Hamming windows). Phase coherence was obtained by means of the *mscohere.m* function from the Signal Processing Toolbox, using 2 s windows with 50% overlap. In Figures 2F and 3D (top panel), for each recording channel we obtained the coherence value at the same frequency as respiration, assessed by the power spectrum of the respiration signal. In Figure 3D (bottom panel), we obtained the coherence value at the same frequency as the theta peak in the LFP power spectrum.

#### Spike-field coupling

Unit activity was obtained by high-pass filtering (>500 Hz). Spike times were defined by setting a threshold above background noise upon visual inspection of individual action potentials. To assess spike-phase coupling, we first filtered the LFP signal within the frequency range of interest based on inspection of power spectral peaks corresponding to RR and theta. Filtering was obtained through a linear finite impulse response filter by means of the *eegfilt.m* function from the EEGLAB toolbox [59; http://sccn.ucsd.edu/eeglab/]. This function applies the filter in the forward direction and subsequently backward to ensure that phase delays are nullified. The phase time series was obtained from the analytical representation of the filtered signal based on the Hilbert transform (*hilbert.m* function). Spike-phase distributions were obtained by associating the spike times with the corresponding instantaneous phases of the filtered signal.

#### Depth profiles of theta and RR

We selected stable periods of theta and RR in which their peak frequencies were separated by >1.0 Hz [25,39]. The LFP signal was then band-pass filtered for isolation of theta and respiration-driven cycles. The precise cutoff frequencies depended on the respiration rate and theta peak frequency, which were determined by inspecting power spectral densities. Recordings from the surface of PAC overlaying the dorsal hippocampus served as the reference for theta [60,61]. Peaks of identified cycles of theta or respiration (assessed from plethysmography, see above) served as triggers for averaging the filtered signals across the 16 electrode positions of the silicon probes (n > 30 waves). To obtain the normalized amplitude shown in Figures 2E and 3C, for each channel we first computed the difference between the peak and trough of its theta- or respiration-triggered average signal, and then divided by the maximal peak – trough difference over channels, so that, for each animal, the maximal amplitude is 1.

#### Intracellular RR amplitude

To estimate the amplitude of intracellular RR cycles (Figure 5C), we first band-passed the membrane potential around the respiration frequency (3 Hz bandwidth) and then collected all voltage values at the peak of individual cycles (notice that the RR-filtered signal oscillates around zero and has no spikes). We also collected the subthreshold membrane potential values at the same timestamps corresponding to the RR peaks. For the latter, spikes were removed by low-pass filtering <10 Hz and any introduced offset from the original signal was manually corrected.

#### Evoked responses

For computing evoked responses to ACC stimulation (Figure 6B), for each LFP channel in the PAC probe we averaged 80-ms time windows triggered by the timestamps of the electrical stimulations (−10 ms to 70 ms; n = 16 to 27 stimulations). Current-source density (CSD) of evoked responses were obtained as previously described [62]. Evoked responses and CSDs were computed individually for each animal and then averaged across animals (n=6 mice).

### Histology

After conclusion of the experiments, the animals were deeply anesthetized with ketamine-xylazine and perfused transcardially with PBS and subsequently with 4% paraformaldehyde (PFA). Brains were carefully dissected, stored in PFA overnight, and coronal sections were cut (50 μm), mounted, and stained with 4,6-Diamidin-2-phenylindol (DAPI) for tracing experiments and electrode localizations. The position of single electrodes and multichannel probes was then verified by fluorescence or light microscopy. Prior to the perfusion, the animals were hyperstimulated using a A365 Stimulus Isolator (World Precision Instruments, Sarasota, FL, USA); applied currents were: 20 μA for 30 s for depth electrodes, 20 μA for 20 s for stimulation electrodes, and 1 mA for 1 s for 16-channel silicon probes.

### Statistics

Data are expressed as mean ± SEM. For comparisons of the proportion of units modulated by theta, RR or both (Figure 4C), we used the Chi-squared test. The statistical significance of spike-phase modulation was assessed through the Rayleigh test for uniformity of phase distribution. We used the Kuiper test to assess the difference between two distributions of the mean theta phase of spiking (Figure 4D). *P* < 0.05 was considered statistically significant.

## Acknowledgments

This work was supported by the Deutsche Forschungsgemeinschaft (SFB 1134/A01; Dr 326/10-1), Bundesministerium für Bildung und Forschung (German-Brazil Cooperation grant No. 01DN12098), the Brazilian National Council for Scientific and Technological Development (CNPq), the Brazilian Coordination for the Improvement of Higher Education Personal (CAPES), and the Alexander von Humboldt Foundation. The funders had no role in study design, data collection and analysis, decision to publish, or preparation of the manuscript.

